# Machine Learning Classification of 53BP1 Foci

**DOI:** 10.1101/2024.02.28.582150

**Authors:** María Xóchitl Benítez-Jones, Sarah Keegan, Sebastian Jamshahi, David Fenyö

## Abstract

**Background:** 53BP1 foci are reflective of DNA double-strand break formation and have been used as radiation markers. Manual focus counting, while prone to bias and time constraints, remains the most accurate mode of detecting 53BP1 foci. Several studies have pursued automated focus detection to replace manual methods. Deep learning, spatial 3D images, and segmentation techniques are main components of the highest performing automated methods. While these approaches have achieved promising results regarding accurate focus detection and cell classification, they are not compatible with time-sensitive large-scale applications due to their demand for long run times, advanced microscopy, and computational resources. Further, segmentation of overlapping foci in 2D images has the potential to represent focus morphologies inaccurately.

**Results:** To overcome these limitations, we developed a novel method to classify 2D fluorescence microscopy images of 53BP1 foci. Our approach consisted of three key features: (1) general 53BP1 focus classes, (2) varied parameter space composed of properties from individual foci and their respective Fourier transform, and (3) widely-available machine learning classifiers. We identified four main focus classes, which consisted of blurred foci and three levels of overlapping foci. Our parameter space for the training focus library, composed of foci formed by fluorescently-tagged BP1-2, showed a wide correlation range between variables which was validated using a publicly-available library of immunostained 53BP1 foci. Random forest achieved one of the highest and most stable performances for binary and multiclass problems, followed by a support vector machine and k-nearest neighbors. Specific metrics impacted the classification of blurred and low overlap foci for both train and test sets.

**Conclusions:** Our method classified 53BP1 foci across separate fluorescent markers, resolutions, and damage-inducing methods, using off-the-shelf machine learning classifiers, a diverse parameter space, and well-defined focus classes.

## Background

By the end of today, endogenous and exogenous agents will have damaged our DNA up to 10^5^ times [1]. Their most severe disruptions will be the double-stranded breaks (DSBs) that they leave behind. Misrepair of DSBs can drive genomic instability, cancer, and aging [2–5]. When a DSB forms, its surrounding chromatin environment changes and cellular signaling events by the DNA Damage Response (DDR) initiate repair [6–8]. Among the earliest signs is the phosphorylation of histone H2AX (γ-H2AX) for long stretches of chromatin surrounding the DSB, known as a focus [9–11]. The chromatin modulator p-53 binding protein-1 (53BP1), another DDR factor, binds to chromatin at the site of damage and also forms higher order structures [12–14]. Detection of γ-H2AX and 53BP1 repair hubs via fluorescence microscopy remains one of the most accurate measures of the total DSB number [15–21]. Methods to quantify foci from fluorescence microscopy images currently consist of manual counting and automated analysis. While manual counting is regarded as the standard, it is subject to large time requirements and biased results [22–25]. Automated methods have increased the efficiency of detection and have the capability of handling multi-channel and multi-dimensional images [23–32].

Resolving overlapping foci that form due to increasing radiation dose remains a challenge in the detection of foci imaged by 2D IF microscopy [25, 32, 33]. Clusters could represent separate foci from distinct planes in the cell or a larger focus from a single plane. Several object segmentation techniques have been explored to address the uncertainty of 2D focus aggregates. Methods differentiating grouped foci have ranged from simple signal thresholding, to complex application of the watershed transform in 3D, convolutional neural networks, and other algorithms [24, 25, 28–32, 34]. Despite the success of these focus segmentation approaches, many come at a high computational cost and overlook the DNA damage complexity information found in clustered foci. While overlapping foci carry uncertainty, segmentation carries the risk of misrepresenting clustered damage sites, which have been observed in cancer cells [35]. Further, previous studies have suggested a link between focus shape and chromatin structure [36–38]. Recently, Vicar *et al* noted a dependence of cell states on the morphology of segmented foci in 3D [25]. For foci imaged in 2D, this relationship remains unexplored. Given that chromatin organization is altered in cancer, exploring the relationship between foci shape and cell state for 2D IF microscopy could have practical implications for clinical and research applications [39, 40].

Focus morphology characterization has applied a mixture of parameter spaces. Detection methods including FoCo, FociCounter, Focinator, and FindFoci use the signal intensity of the focus as their main parameter [23, 26, 28, 34]. While the focus intensity as the sole parameter is a simple feature, it is prone to batch effects such as signal variations across samples and experiments. Some detection methods include additional measures. FocAn and FociPicker measure the signal statistics and the geometric properties of the foci, respectively [27, 31]. Various machine learning methods have been implemented for focus detection. Hohmann *et al*‘s identification and classification of foci based on their quality used a features space containing filters [30]. Deep learning approaches, such as FociNet, DeepFoci, and FociRad, have implemented convolutional neural networks to detect foci [24, 25, 32]. SMLM studies that characterized foci have implemented cluster analysis and measured the topology of focus-forming proteins, for example 53BP1 and γ-H2AX, which include variables such as the number of components and the number of holes [37, 41, 42]. While deep learning approaches have produced some of the best performing detectors which could be efficiently implemented once trained, and SMLM has characterized foci at the nanoscale, these methods currently require retraining for implementation and necessitate resources that may not be available to all laboratories. Further, additional morphological properties, such as frequency domain measurements, remain unexplored.

The total number of foci detected serves as a distinguishing marker to classify cells based on the amount of DNA damage present in a given cell. Focus detection methods have classified cells based on the focus count alone. Chen *et al* distinguished cells that had varying levels of DNA damage by using convolutional neural networks to detect the number of foci in each cell [24]. Although Hohmann *et al* did not classify cells, they classified foci to determine true signals, which improved focus detection quality, as mentioned earlier [30]. Deep learning remains one of the best methods for focus detection, leading to better cell classification. However, Vicar *et al* observed that to distinguish cells based on disease and DNA repair times, parameters beyond focus count were required [25]. Even while achieving the highest Dice score compared to other focus detection methods by using deep learning, Vicar *et al* were only able to resolve cell types by measuring focus properties. A machine learning approach that improves focus detection quality and classifies cells based on focus properties using readily-available resources has been unexplored. In addition to deep learning, some of the best focus detection methods rely on 3D datasets [25, 27, 29, 31]. Many of these focus detection methods require large and complex datasets, specific microscopy hardware and software, storage space, processing power and time in training. While a trained deep learning model could detect foci efficiently, current deep learning focus detection methods require training for implementation. To implement focus quantification from scratch, computationally efficient methods are necessary for clinical applications like dose estimation [43, 44].

Here, we present a 53BP1 focus classification method using straightforward machine learning approaches based on a diverse parameter space. We have implemented two types of fluorescence microscopy, suggesting that our method has the potential to generalize across different fluorescence microscopy methods to quantify 53BP1 foci. Foci from two datasets with varying levels of DNA damage are characterized via a novel focus class framework and parameter space measuring geometric and statistical properties of individual foci in their spatial and frequency domains. Our classes provide a framework applicable across methods that induce random DSBs, 53BP1 markers, and fluorescence microscopy techniques. Focus quality was determined by differentiating blurred foci from other signal classes. Application of this method can range from areas of DNA repair research to clinical applications involving foci immunohistochemical assays.

## Methods

### Training dataset

#### Cell culture and sample preparation

Our cellular system consisted of female human bone osteosarcoma U2OS cells (ATCC HTB-96) which expressed BP1-2 (53BP1 aa 1220-1711) fused to EGFP and different halo-tagged DNA repair proteins (data not included). This marker identified 53BP1 without its functional domains [45]. Cell culturing medium consisted of DMEM with 100 U/mL Penicillin-Streptomycin (Gibco 15140122) and 10% fetal bovine serum (Gemini Bio Products 100106). Cells were incubated in a chamber at 37°C and 5% CO2. To seed cells, they were trypsinized and plated in glass bottom dishes (ibidi 81218). Cells were synchronized in G1/G0 via a 48-hour incubation in medium without FBS but with Pen-Strep. Samples treated with phosphorylation inhibitors were incubated for 15 minutes. DSBs were generated through a 10-minute neocarzinostatin incubation (NCS, Sigma-Aldrich, 100 ng/mL). Cells were PBS-washed three times, incubated (15 minutes) and imaged in FluoroBrite^TM^ DMEM (Gibco A1896701).

#### Live cell imaging

Imaging was conducted on a custom-built inverted Rapid Automated Modular Microscope (RAMM) System (ASI) with an XYZ automated stage (ASI PZ-2000 FT) and a 150X objective with a 1.45 NA (UAPON150XOTIRF Olympus). Samples were illuminated with 639 nm (MRL-FN-19052399), 473 nm (Laserglow Technologies LRS-0473-PFF-00200-05), and 405 nm (MDL-III-405) lasers; each passed through their respective clean-up filters (ThorLabs) before reaching the sample. Emissions traveled through a 2X tube lens and a two-way image splitter (Photometrics OptoSplit II), which included a dichroic mirror (Semrock 660 Di), band pass filters (605/75, 706/95, 531/40), long pass filters (647, 532, 561), and a notch filter (405/473/532/639). A sCMOS camera (Photometrics Prime BSI) captured the separated emissions. A total of 2000 frames were acquired for each region of interest at 33 Hz. Approximately two nuclei were imaged per region of interest.

#### Image processing and focus detection

Images from both channels were collected and mapped. ROIs of single nuclei were manually selected (Fig. 1A) using the maximum projection of the raw images. ROIs of individual nuclei were extracted from the raw EGFP-channel data and processed to improve the signal to noise ratio (Fig. 1B). Standard image processing steps were applied to three evenly-spaced frames in the EGFP channel ROI stack. Contrast-limited adaptive histogram equalization via the createCLAHE function from OpenCV library increased contrast. Hot pixels were removed by subtracting the histogram-equalized median-blurred image (medianBlur function from OpenCV library with a 3x3 kernel) from the histogram-equalized image. Gaussian blurring was employed with the GaussianBlur function from OpenCV library with a 7x7 kernel and a 1x1 kernel for the train and test set images, respectively. To implement background subtraction via k-means clustering (Fig. 1B), images were reshaped into a column vector and inputted into the kmeans function from the OpenCV library with k = 4, a maximum iteration of 10, an epsilon of 1, and a *KMEANS_RANDOM_CENTERS* flag. Centers were sorted and the image was thresholded using the third largest center. Pre-processed images were mean-projected (Numpy’s mean function), thresholded with Otsu’s method (scikit-image’s threshold_otsu function), and the remaining objects were detected by connected component labeling (Mahotas’ label function) (Fig. 1C). Focus size was thresholded using minimum and maximum total pixel counts.

**Figure 1:**
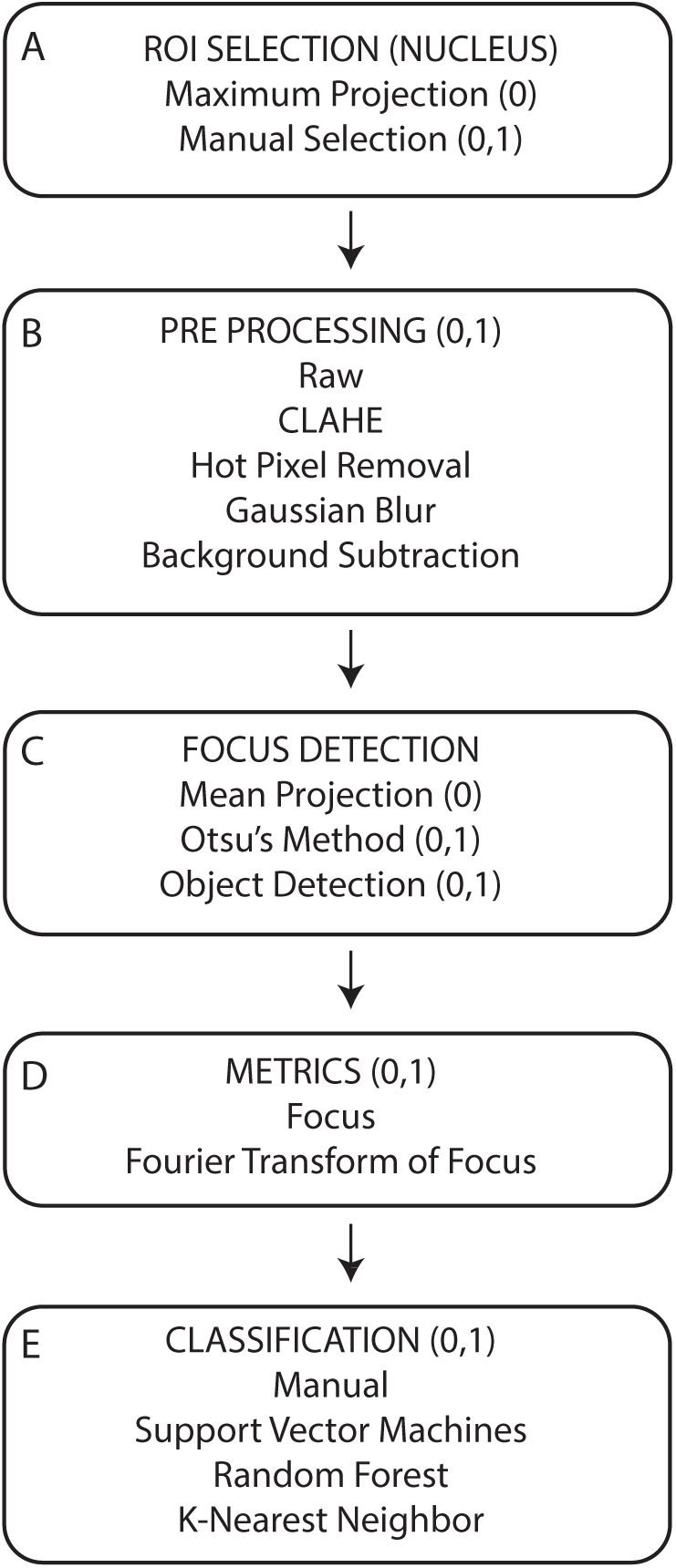
Analysis pipeline. Training set steps are labelled (0) and testing set steps are labelled (1). A) A manually selected ROI containing the nucleus of interest was B) processed to enhance the signal-to-noise via contrast-limited adaptive histogram equalization, hot pixel removal, gaussian blurring, and background subtraction with K-means clustering. C) Foci were detected within each nucleus ROI using Otsu’s thresholding and connected component labeling. D) Properties for each detected focus and their Fourier transform magnitude were measured. All detected foci were filtered using the detected focus ROI’s total pixel count. E) Manual and automated classification of foci Vector Machines, Random Forest and K-Nearest Neighbors.

### Testing dataset

A subset of open source high content microscopy data performed by Doil *et al* available on Image Data Resource (IDR, https://idr.openmicroscopy.org/webclient/?show=plate-4501) constituted our testing dataset [46, 47]. The screens contained U2OS cells seeded onto 5 ng siRNA spots. Each screen included 384 siRNA spots and each spot was imaged on a wide-field microscope. Cells were fixed, permeabilized and endogenous 53BP1 was immunostained with Alexa Fluor 488. Details on siRNA arrays, image acquisition, and additional methods implemented by Doil *et al* have been previously described [46]. Single-nucleus ROIs were manually selected and rescaled by matching the training library’s focus ROI pixel size median. ROIs were pre-processed (Fig. 1B) and foci were detected as previously described (Fig. 1C), with the exception of including a mean projection as spot images acquired by Doil *et al* consisted of a single time point captured at 33 Hz.

### Parameter space

Focus morphologies found in the training and testing datasets were classified via the parameter space created by properties measured for each focus ROI and its Fourier transform spectrum (Fig. 1D, Table 2). Statistics of an ROI’s intensity, such as its maximum, mean, median, minimum, sum standard deviation, and variance were measured. The total pixel number and the Euler number of the ROI was also included. Geometric properties included area, compactness (Polsby-Popper and Schartzberg), eccentricity, major axis length, minor axis length, orientation, perimeter, and solidity. Histogram comparisons projected an ROI ‘s intensity onto x and y axes to obtain two histograms and compared them by way of distance (Alternative Chi Squared, Bhattacharyya, Chi Squared, Hellinger), the Kullback-Leibler divergence, the histograms’ correlation and intersection. A perceptual metric [48] and the variance of the Laplacian were used to quantify blur. The latter detected the edges of the object of interest by calculating the Laplacian of its gaussian blur and measured its variance.

### Class definition and manual classification

The imaging plane within a living cell provides a limited view of foci, where some foci are resolved and other foci are blurred. Different forms of resolved foci observed in the training set were categorized into three classes according to their degree of overlap: low, moderate, and high overlap (Fig. 2). We therefore defined the following classes: background (a), misdetection (b), blurred foci (0), and foci with low (1), high (2), and moderate (3) overlap levels. Since our study explored focus classification rather than focus detection optimization, we considered classes a and b as artifacts which were not included in the focus training library. Because blurred foci in class 0 consisted of foci that were outside of the imaging plane they were regarded as noise. Classes 1-3 were defined as signal within the imaging plane. Class 1 consisted of circular foci that were relatively symmetric compared to classes 2 and 3. Multiple overlapping foci with an elongated signal that broke the symmetry visible in class 1 were defined as class 2. Class 3 was composed of foci with moderate overlap. Following detection and metric measurement, foci from both libraries were manually classified (Fig. 1E). Unclear cases were determined with a flowchart (Fig. S1).

**Figure 2:**
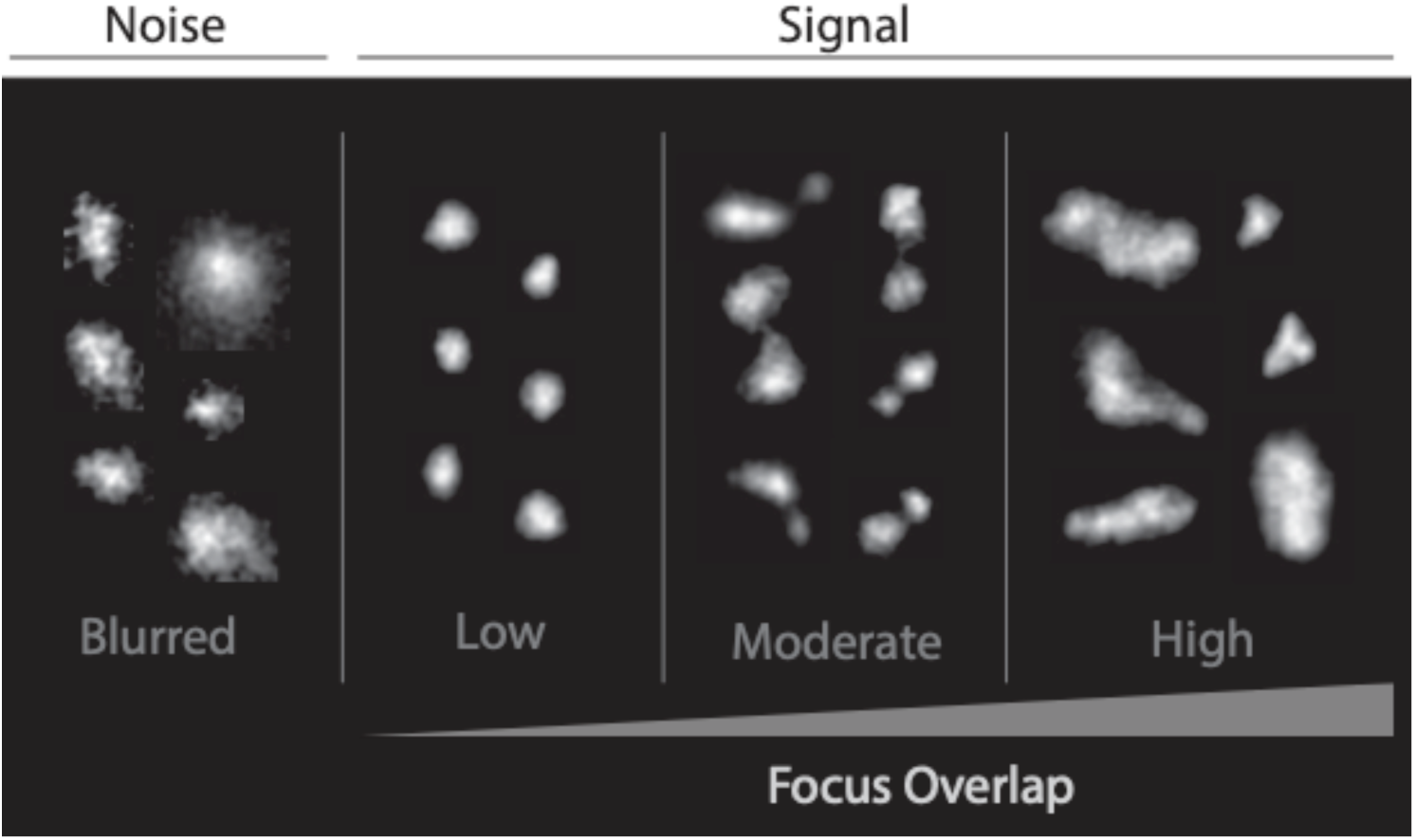
Focus class framework. Training set foci were divided into foci that exhibited ill-defined edges (noise) and well-defined edges (signal). The signal class contained subclasses of foci with low, moderate and high levels of overlap. The noise class was synonymous with blurred foci.

### Training and testing

The training library was composed of a total of 4507 individually detected foci from 92 nuclei, their measured properties and class (Table 1). Foci were normalized with respect to their ROI’s total pixel number and properties were normalized by their mean and scaled. The library was split into sub-train and sub-test sets. Machine learning classifiers included Random Forest, Support Vector Machines, and k-Nearest Neighbors (Fig. 1E). Classifications were performed as both binary and multiclass problems. We implemented Scikit-learn’s *OneVsRestClassifier* to evaluate binary problems via ROCs and their AUC. Here, the multiclass problem was reduced to several binary classification problems. One class was labeled as the positive class, while the rest were labeled as the negative class. Binary classification was iterated for all four classes. Ten-fold cross-validated weighted metrics (F1 score, precision, and recall) evaluated multiclass performance for all classifiers and classes. Given that focus symmetry and blur measurements have been unexplored, we investigated the classifier dependence on these metrics. Properties that separated the distributions of blurred/low overlap classes (Fig. S6A, Table 2) and low/high overlap classes (Fig. S6B, Table 2), were identified as blur and symmetry metrics, respectively. We compared the classification of low vs high overlapping foci in the presence and absence of the symmetry-measuring properties from foci and their Fourier transform. A similar comparison was performed for the blur metrics in distinguishing the low overlap and blurred classes. The number of detected foci in the test dataset totaled to 1165 from 363 nuclei. Just as with the training library, the testing library included focus properties and classes (Table 1). Previously-trained classifiers were evaluated with the test set for the binary and multiclass problems, as well as the metric dependence experiments outlined above.

**Table 1:** Detected focus counts. Total number of manually classified foci (signal and noise classes) for the train and test datasets.

|  | Low Overlap | Moderate Overlap | High Overlap | Blur | Total |
| --- | --- | --- | --- | --- | --- |
| **Train** | 3109 | 200 | 319 | 879 | 4507 |
| **Test** | 847 | 54 | 70 | 194 | 1165 |

**Table 2:** Measured properties for each focus. HC: Histogram comparison, FT: Fourier Transform, VL: Variance of Laplacian, GB: Gaussian Blur, s: symmetry-measuring properties, b: blur-measuring properties.

| Label | Property Name | Label | Property Name |
| --- | --- | --- | --- |
| 0 | Orientation | 24 | FT Schwartzberg Compactness |
| 1 | Area (s, b) | 25 | FT VL |
| 2 | Perimeter (s, b) | 26 | FT Perceptual Blur |
| 3 | Major Axis Length (s, b) | 27 | FT Shannon Entropy (s, b) |
| 4 | Minor Axis Length (s, b) | 28 | FT Mean |
| 5 | Eccentricity | 29 | FT Median |
| 6 | Correlation HC | 30 | FT Sum (b) |
| 7 | Chi Squared HC | 31 | FT Standard Deviation |
| 8 | Intersection HC | 32 | FT Variance |
| 9 | Bhattacharyya HC | 33 | FT Maximum |
| 10 | Alternative Chi Squared HC | 34 | FT Minimum |
| 11 | Hellinger HC | 35 | VL |
| 12 | Kullback-Leibler Divergence HC | 36 | VL GB 3x3 Kernel |
| 13 | Euler Number | 37 | VL GB 5x5 Kernel |
| 14 | Solidity (s) | 38 | VL GB 7x7 Kernel |
| 15 | FT Orientation | 39 | VL GB 9x9 Kernel |
| 16 | FT Area | 40 | Perceptual Blur |
| 17 | FT Perimeter | 41 | Shannon Entropy (s) |
| 18 | FT Major Axis Length | 42 | Focus Mean |
| 19 | FT Minor Axis Length | 43 | Focus Median |
| 20 | FT Eccentricity | 44 | Focus Sum |
| 21 | FT Euler Number | 45 | Focus Standard Deviation |
| 22 | FT Solidity | 46 | Focus Variance |
| 23 | FT Polsby-Popper Compactness | 47 | Polsby-Popper Compactness |
|  |  | 48 | Schwartzberg Compactness |

## Results

We aimed to establish a focus classification system that was concise and generalized well across experimental conditions, and build models for automatically classifying foci. The signal and noise were identified as the two main classes (Fig. 2). The signal class was split into subclasses of low, moderate, and high degrees of overlapping foci. Blurred foci with ill-defined borders, compared to the well-defined borders of foci in the signal classes, were assigned to the noise class. Once foci were detected, they were manually labeled according to the established class framework; one class per focus. Our approach to building an extensive parameter space, a key element of our method, included measuring a wide array of fifty-two morphological and statistical properties of the labeled foci and their FT. To ensure the utility of our parameter space and the generality of our classes, we selected two datasets that differed with respect to DNA damage induction, microscopy method, 53BP1-labeling technique, resolution, and cell treatment. Our training dataset consisted of live cell imaging data of BP1-2 foci (53BP1 without its functional domains) tagged to EGFP. We tested our trained classifiers with a publicly-available high content microscopy screen dataset of endogenous 53BP1 foci immunostained with Alexa 488.

### Establishment of Focus Parameter Space

To assess the dependence between properties in our parameter space we used heatmaps to visualize their Pearson correlation coefficients (Fig.3, Table 2). For the training dataset, the focus geometric properties, perceptual blur and sum, as well as the Fourier transform of the focus’ geometric properties, Shannon entropy, sum, maximum (properties 1-4, 16-19, 27, 30, 33, 40, 44) showed a moderate to strong negative correlation (<-0.40) with the solidity, Polsby-Popper and Schartzberg compactness of the Fourier transform of the focus as well as the variance of the Laplacian with different-sized gaussian blur kernels, both compactness metrics, mean and median of the focus (properties 22-24, 36-39, 42, 43, 47, 48). A strong positive correlation (>0. 70) was observed between properties 1-4, 16-19, 27, 30, 33, 40, and 44. Smaller groups exhibited strong positive correlation. The focus mean, median, and both compactness metrics (properties 42,43,47,48) scored correlation values greater than 0.60. The Fourier transform variance of the Laplacian, standard deviation, and variance of the focus (properties 25,31,32) were strongly correlated (>0.80). Properties that did not exhibit linear relationships included the eccentricity of the focus FT (property 20) and focus orientation (property 0), scoring correlation values under 0.1 for several properties.The histogram comparison via correlation (property 6) demonstrated moderate negative correlation (-0.50) with a variety of properties including the focus eccentricity, histogram comparison via standard and alternative Chi-Squared distance (properties 5, 7, 10), and a strong negative correlation (-0.80) with the histogram comparison via Bhattacharrya and Hellinger distances (properties 9, 11). The variety of correlation between variables indicated that our parameter space was a diverse input for machine learning classifiers.

**Figure 3:**
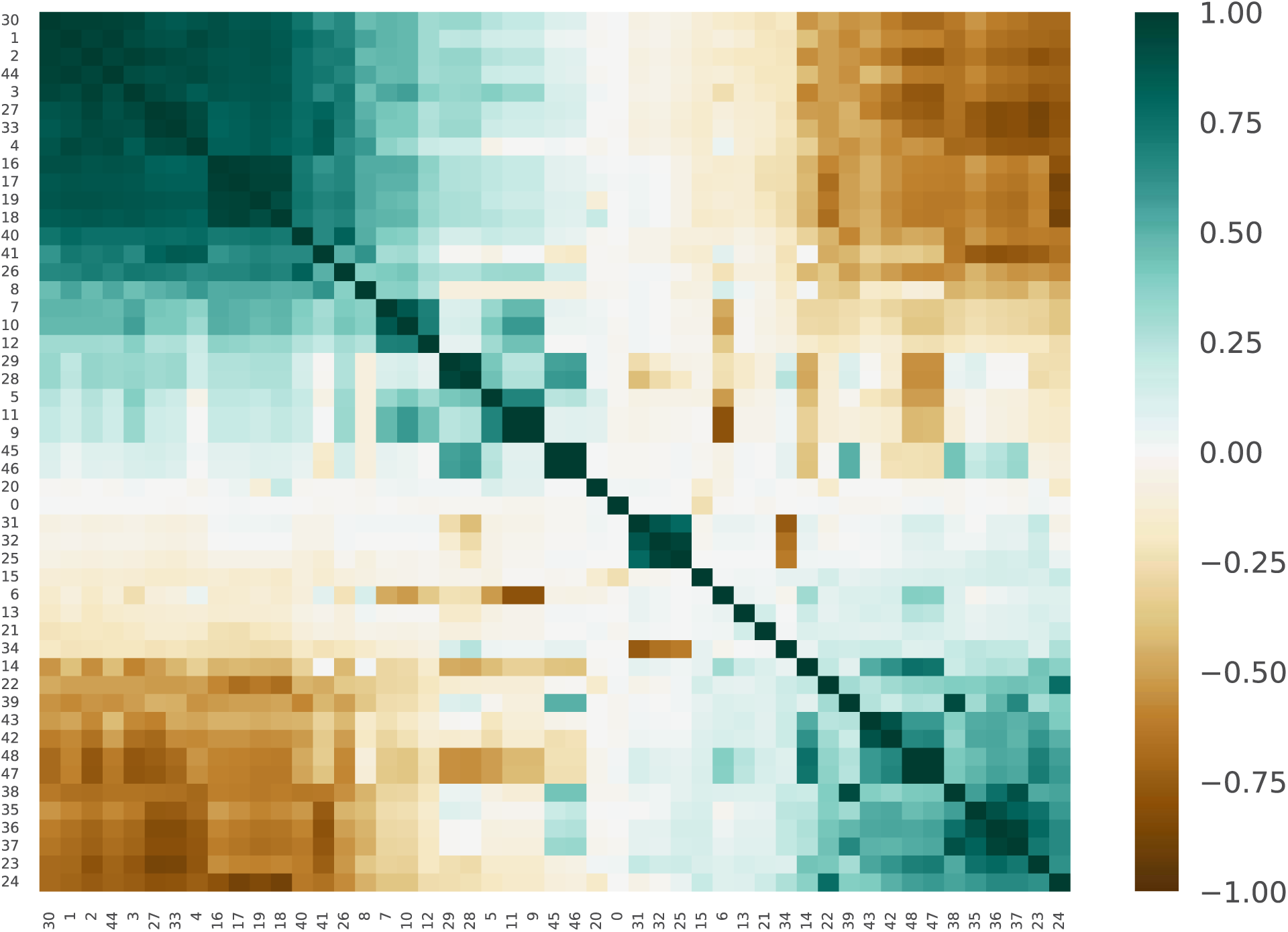
Parameter space. Pearson correlation coefficient of the properties was measured for detected foci in the symmetric, asymmetric, midway, and blur classes for the train dataset and visualized via heatmaps. Measured properties consisted of geometric, blur, symmetry, and frequency spectrum measurements.

The correlation heatmaps of the properties measured for the test foci and their FT were also plotted (Fig. S2A, Table 2). The test dataset demonstrated a strong to moderate negative correlation (<-0.40) between the same properties observed for the train set plus the focus’ Shannon entropy (property 41), with the exception of the mean and median (properties 42, 43) of the focus, which demonstrated a moderate to weak positive correlation (<0.40). A similar strong positive correlation (>0.70) was also observed between properties 1-4, 16-19, 27, 30, 33, 40, 41, and 44. The smaller groups echoed the higher positive correlation in the train set. One small group of properties, the Fourier transform variance of the Laplacian, standard deviation, and variance of the focus (properties 25, 31, 32), with moderate to strong positive correlation (>0.50) was validated in the test set. A similar variety of properties observed in the train set demonstrated weak negative (>-0.10) and positive correlations (<0.10) with the focus orientation (property 0) and eccentricity of the focus FT (property 20), respectively. Just as with the train set, moderate negative correlation (<-0.50) was observed between the histogram comparison via correlation (property 6) and a variety of properties including the focus eccentricity, histogram comparison via standard and alternative Chi-Squared distance (properties 5, 7, 10), and a strong negative correlation (-0.70) well as the histogram comparison via Bhattacharrya and Hellinger distances (properties 9, 11).

Property continuity between datasets was observed in a test *vs.* train correlation value scatter plot (Fig. S2B, Table 2). Thirty-seven properties exhibited consistent correlation values between the testing screen and the training live cell data. The remaining twelve training properties that did not fully translate to the test foci included the focus mean, median, variance, standard deviation, Shannon entropy, variance of Laplacian with (9x9 kernel) and without gaussian blur, Polsby-Popper and Schartzberg compactness, perceptual blur, and the histogram comparison via Chi-Squared distance and Kullback-Leibler divergence (properties 7, 12, 35, 39-43, 45-48). These results supported that foci induced, imaged (live-cell *vs.* immunofluorescence), and labeled using two distinct reporters (endogenous 53BP1 *vs.* BP1-2-EGFP), in two separate labs, and at two different times, shared similar properties.

### Focus Classes Distinguished Across Datasets

After exploring our parameter space, we applied random forest (RF), support vector machines (SVM), and k-nearest neighbors (KNN) to classify our training set. The binary problems we implemented evaluated the classification of one class against the rest of the classes; the latter were considered a single class (Fig. S3). The moderate overlap class was the easiest to classify, where all classifiers scored an AUC of 0.93 or higher. Also well-classified were the low and high overlap classes, where an AUC of 0.90 or higher was observed for all classifiers. The blurred class exhibited the lowest scores, where all classifiers scored an AUC between 0.85 and 90. SVM and RF held the highest AUCs for all classes; their ROCs were similar, with the exception of the high overlap class. Random forest held the highest true positive (TPR) to lowest false positive rate (FPR) ratio early on when classifying the high overlap class, until SVM surpassed RF after an FPR of 0.20. KNN maintained a score above 0.84 for all classes. Classifier performance was further assessed for multiclass problems, a case where all classes were included. Each classifier was evaluated via 10-fold cross validated recall, F1 score, and precision metrics. RF scored the highest recall and F1 medians, which echoed the higher performances observed in the binary problems (Fig. S4). SVM scored the highest precision median, albeit with an inconsistent performance regarding the remaining evaluation metrics. KNN mostly scored the lowest medians, with the exception of its recall median. As classifiers for the training library, RF and SVM were neck-and-neck as the best, with KNN not falling far behind.

Similar to the training set, the classifiers were evaluated as binary problems (Fig. 4) on the test set. In this dataset, the easiest class to distinguish for all classes was the moderate overlap class, all classifiers having scored AUCs of 0.90 or higher. As with the training set, the blurred class was not classified as easily as the other classes. The AUC scores ranged from 0.75 to 0.80 for the blurred class, 0.11 lower than the training set on average. SVM maintained the highest AUCs and curves for the moderate and high overlap classes. RF performed the best when classifying the low overlap and blurred classes. In general, the signal classes were easiest to classify for both the training and test set, while the noise class was the most difficult to distinguish. Lack of continuity regarding the definition of a blurred focus in both datasets most likely influenced a lower performance. Shifts in the results were observed when the classifiers were tested in the multiclass problems, albeit with an analogous overall pattern (Fig. S4). Although RF did not score the highest F1 score, all of the classifiers’ precision and recall scores followed similar trends observed in the training set. Among the higher performing classifiers remained RF, which was mirrored by KNN, while SVM sustained a varied performance. The top three features for RF were the focus sum, histogram comparison via intersection, and minor axis length (Fig. S5). All scores for the multiclass problems were higher for the training set compared to the test set.

**Figure 4:**
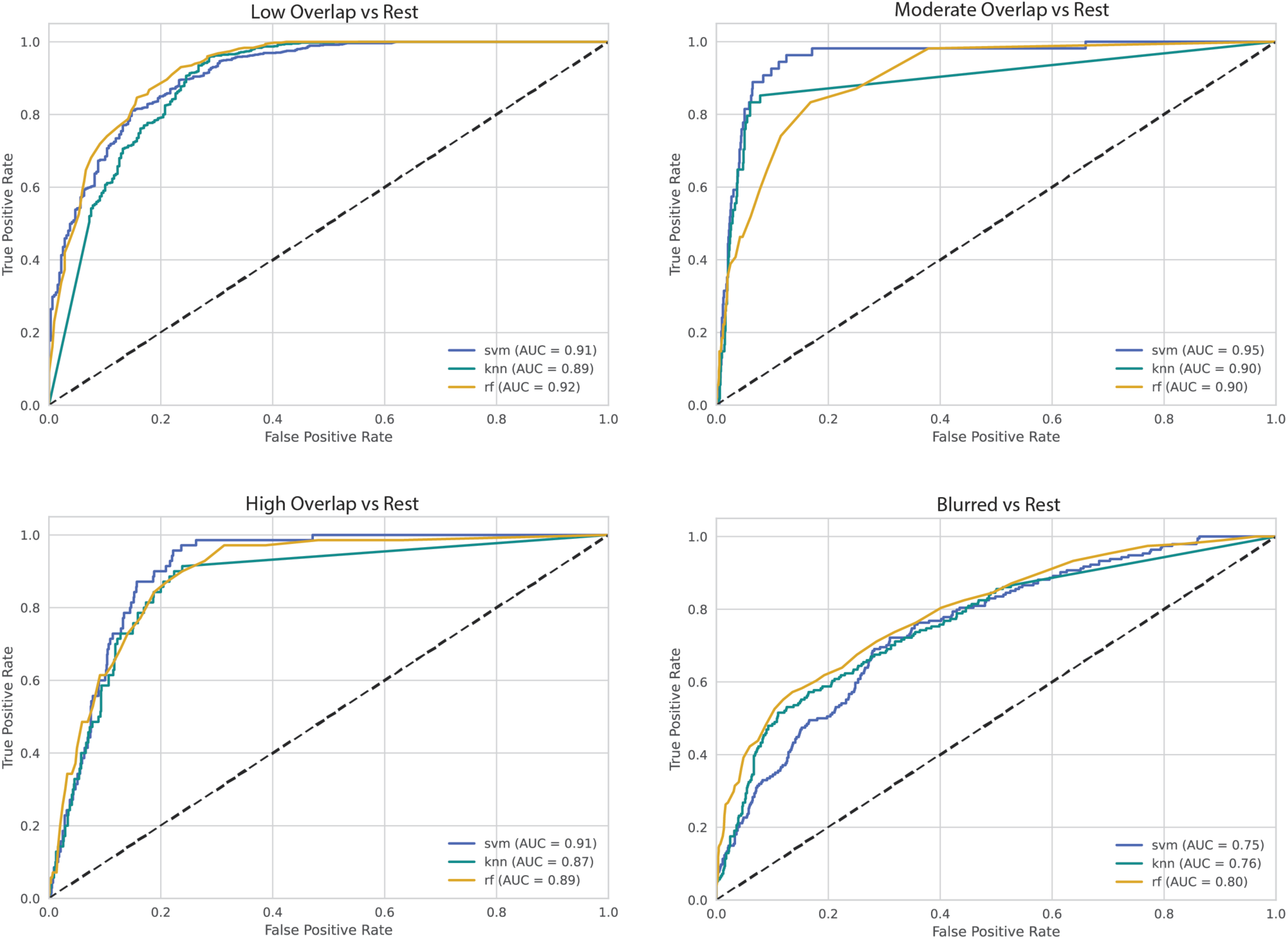
Classifier evaluation. Binary problems via One-vs-Rest classification measured the performance of RF, SVM, and KNN for the signal and noise classes using the test sets. The ROC curves and AUC scores evaluate the classifiers’ performance for each class.

### Effects of Symmetry and Blur Metrics on Focus Differentiation

Dependence of focus classification on symmetry and blur metrics was evaluated by a 10-fold cross validation of the recall, F1 score and precision for RF, SVM, and KNN using a subset lacking symmetry or blur metrics to compare against the full set of metrics. Distinction between the low and high overlap classes was observed to evaluate the effect of the symmetry metrics (Fig. 5A, Fig. S6A). We observed that the distribution for RF and KNN presented the highest median recall and F1 scores for the train (Fig. S7A) and test (Fig. 5A) datasets. After the symmetry properties were dropped, there was minimal to negligible effect on the classifiers’ recall and F1 medians, with the exception of SVM, which scored lower recall and F1 medians without the symmetry metrics. All three classifiers scored similar precision medians for the train and test sets; SVM only displayed a notable drop without the symmetry metrics for the train and test sets. RF and KNN mostly scored the highest metric medians and were not dependent on the identified symmetry metrics for low *vs.* high overlap focus classification. SVM scored lower evaluation metric medians overall, and was sensitive to the drop in the defined symmetry metrics.

**Figure 5:**
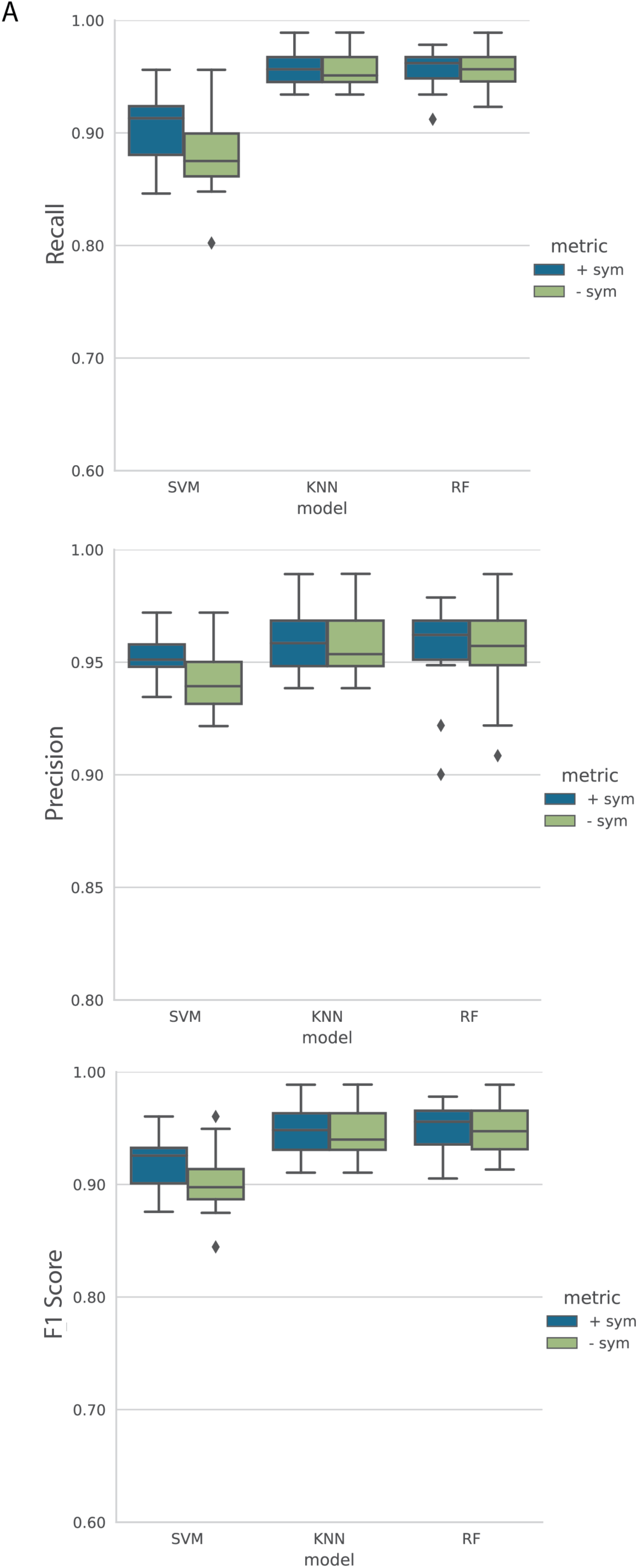
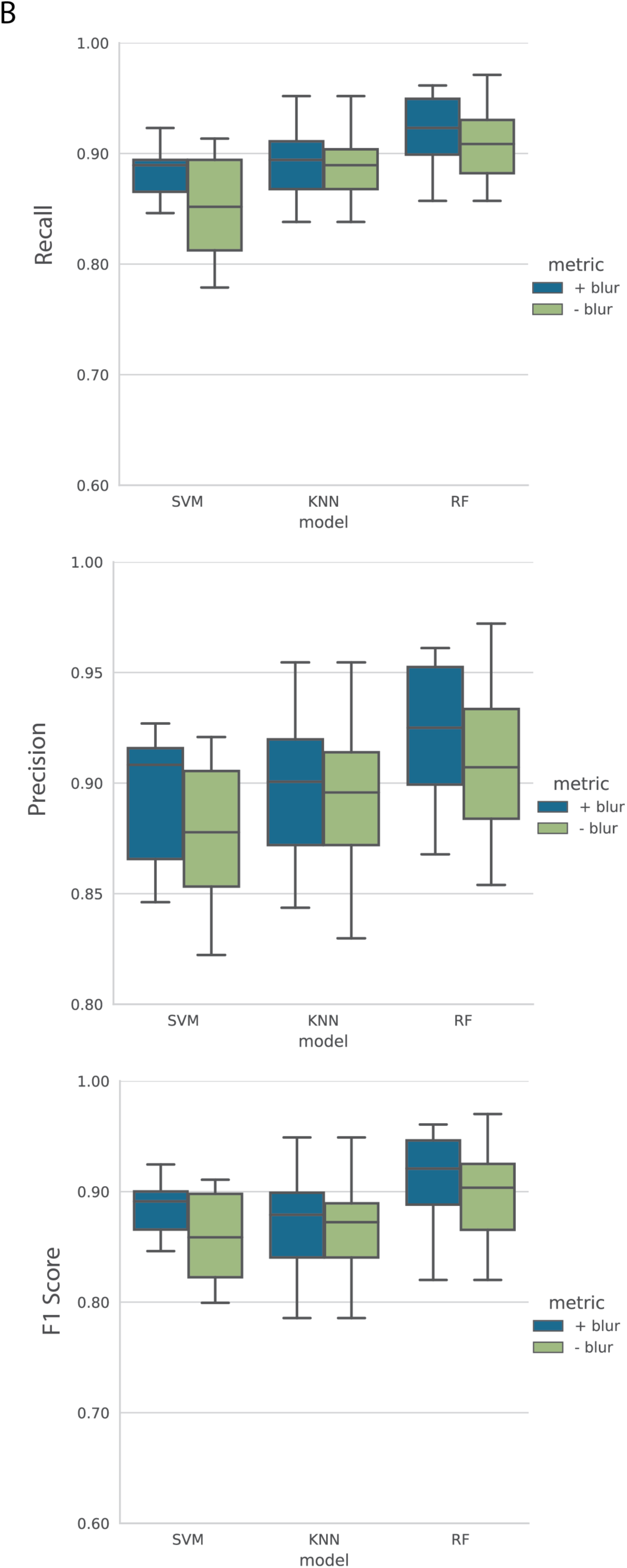
Effect of metrics on classification. Boxplots of the classifiers’ 10-fold cross-validated recall, precision, and F1 scores in cases where key symmetry or blur metrics were included (+, blue) and removed (-, green) to observe the effect on the differentiation between classes. A) Classification of low overlap vs high overlap class in the presence and absence of symmetry metrics for the test set. B) Distinction between blur and low overlap class in the presence and absence of blur metrics for the test set.

Many images include blurred foci, but these are not usually separated from foci that are in the focal plane. Distinguishing these blurred foci would decrease the uncertainty in focus type. In the parameter space, we observed that there were properties that distinguished the blurred class from the low overlap class (Fig. S5B). We compared the classification of blurred foci in the presence and absence of the blur properties of the foci and the Fourier transform of the foci for the train and test sets. Random forest consistently exhibited high scores for the train (Fig. S7B) and test (Fig. 5B) datasets, which dropped without the blur metrics. Support vector machines scored some of the highest medians for the train set, but did not maintain the highest scores for the test set. Most k-nearest neighbors’ medians were among the lowest with the blur metrics for both datasets. While a decrease in KNN’s performance in its differentiation of blurred foci and low overlap foci was observed without the blur metrics, this difference was minimal in the test set.

## Discussion

Automation is well under way to replace manual focus detection. In contrast, automation of focus classification is largely unexplored. Manual differentiation of foci requires the definition of general classes and their application across samples and experiments, which is prone to bias and time constraints. Machine learning methods can clearly and quickly define and classify different types of foci, decreasing these limitations. Attempts at detecting and measuring the properties of these foci have been overly simple or overly complex [25, 27]. FociPicker measured the focus intensity, area, volume, and coordinates, and applied a thresholding algorithm to separate overlapping foci [27]. In contrast, DeepFoci used deep learning to detect foci and measured morphological properties of 3D foci in addition to the intensity value [25]. Our results demonstrated that given an extensive parameter space and general classes, support vector machines, random forest, and k-nearest neighbors differentiated focus types across experiments involving different cell markers, imaging methods with different spatial resolutions, and DSB generation techniques. These methods are not as time-consuming as deep learning. For example, the total computational training time for FociRad’s deep learning focus detection was over 37 hours [32]. Our method’s total computational training and testing time classified foci within an hour (data not shown). Microscopy, storage space, and computational power are additional barriers to entry when using deep learning. Many of the datasets employed by deep learning detection methods used 3D image stacks [25, 32]. A single 3D multi-channel image from DeepFoci measured roughly 8 MB. The high content microscopy images from Doil *et al* that composed our test set were approximately 700 KB per image [25, 46]. The computational tools employed by Wanotayan *et al*’s FociRad, a deep learning detection model, were an Intel 1151Core i5-9600K processor, GeForce RTX 2070 GPU, and 250 GB of RAM [32]. Although our computational tools were also advanced (Intel Core i7-12700 processor, NVIDIA GeForce RTX 3080 GPU, and 32 GB of RAM), other platforms, such as CellProfiler Analyst, could apply random forest, or other classifiers, to distinguish between our defined focus classes using the features in our parameter space [49]. Measuring a variety of focus properties and employing simpler machine learning methods for 2D immunofluorescent images is an underexplored middle ground between thresholding algorithms and deep learning.

The number of DNA damage foci per cell is a popular measure to distinguish cellular states with respect to the amount of damage induced. Measurement of DNA damage is implemented from diagnostics to basic DNA repair research. Many of these fields necessitate quick assays that assess damage at scale [50]. Capturing immunofluorescent images of foci has been among the most accurate DNA damage-measuring assays, which can be productively automated to increase its capacity and speed [51]. Automation of focus counting assays, such as the rapid automated biodosimetry tool (RABiT), have been developed for radiological occurrences and beyond [52]. Focus counting remains the analysis of choice for these images. While it is a straightforward measure of DNA damage, it misses information contained in the focus morphology. Focus counts are typically composed of circular foci, which is not a comprehensive representation of the focus landscape present in the nucleus. Overlapping foci occur due to an increase of number and/or size with an increase in radiation. Exclusion or inclusion of overlapping foci as circular foci has led to miscounts in focus enumeration [32, 53]. Further, some focus detection methods include foci that are out of the focus plane, where the risk of classifying multiple foci as a single focus is high due to the uncertainty brought on by blur [30, 32]. To our knowledge, ours is the first focus classification method that measures blur and can distinguish blurred foci from high SNR foci, as well as differentiating between discrete and varying levels of overlapping foci. Well-defined and generalizable focus classes could serve as a replacement for focus counts as radiation damage markers. An alternative to the rate of circular foci per cell is the rate of focus classes per cell. The focus class framework developed here could classify foci in cell samples and determine the amount of low, moderate, and high overlapping foci per cell. The rate of blurred foci per cell serves as a quality indicator of the data. Given these rates, inference of a cell’s state in regards to the amount and complexity of induced damage can be determined. With this framework, focus counting has the potential to classify cells [54]. While experiments on cell classification via focus class rates remain to be explored, previous observations on the requirement of focus morphological measurements for cell class distinction by Vicar et al. are promising [25]. Calibration curves for radiation dose estimation require the focus count per cell[55–57]. The focus count is plotted against the dose and fitted using linear regression to quantify the relationship between dose and the amount of damage induced. As mentioned above, encounters with overlapping foci at high doses limit the functionality of dose-response curves generated with focus counts. However, hidden opportunities lie in the morphology of overlapping foci. The focus class framework could be implemented by plotting several dose-response curves with respect to the number of focus classes, i.e. the number of high overlapping foci per cell per dose, the number of low overlapping foci per cell per dose, etc. Dose-response curves have not been generated with respect to overlapping focus classes and remain to be explored. However, since high overlap foci are suggestive of more complex DNA damage and higher dose, formation of focus class-specific dose-response curves is within reach. The next steps in our method would involve improving the detection of the blurred foci via further analysis of the frequency domains and properties that more accurately measure blur as well as metrics that have lower noise sensitivity and measure the geometry of foci. The signal classes were easily distinguishable by all classifiers, and could be used to obtain new dose response curves.

Various focus-quantifying metrics foci were explored in our method. The observed parameter space contained a range of correlation clusters between properties (Fig. 3, Table 2). These ranged between strong, moderate, and weak positive and negative correlations. Properties exhibiting positive correlation clusters included geometric properties, pixel statistics, and histogram comparisons of the focus, as well as geometric, pixel statistics, and Shannon entropy of the magnitude of the FT of the focus. Negatively correlated clusters mostly involved the variance of the Laplacian, a few geometric properties and histogram comparison, and the topological measure of the focus, in addition to a few geometry properties, variance of the Laplacian, pixel statistics, and the topological measure of the magnitude of the FT of the foci. Future dimensionality reduction of the metric clusters would simplify the parameter space and indicate which focus-characterizing properties would carry the most weight within each cluster. Inclusion of properties that measure focus symmetry and blur was new territory that we explored (Fig. 5). By eye, poorly-defined edges indicated a blurred focus and smooth edges indicated a signal focus. In distinguishing between the low and high overlap foci, we hypothesized that the histogram comparisons and geometric properties would play key roles. While the properties that separated these two classes (Fig. S6A) relied mostly on the geometric properties of the focus and magnitude of the FT of the focus, there was a minimal to negligible effect on the classification of these classes without the geometry metrics (Fig. 5A). We hypothesized that the classification of the blur class relied heavily on the variance of the Laplacian. To our surprise, properties that separated the distributions of the blurred and low overlap foci (Fig. S6B) largely included focus geometric properties, but did not include its variance of the Laplacian. These properties were determined by their ability to separate class distributions. Assigning these properties based on their Pearson correlation coefficients would be another avenue we could implement to improve the symmetry and blur metric definitions. Concerning classifier performance, all classifiers performed similarly, with a few distinctions. RF demonstrated the most stable scores across datasets for the binary and multiclass problems. The highest performance in the binary problems was achieved by SVM, albeit with an inconsistency exhibited in the multiclass problems. Although KNN mostly scored lower than RF and SVM, it performed consistently in the binary and multiclass problems. Given that all three classifiers performed relatively well, our next steps would implement majority voting to improve the classification performance.

Our study contained limitations. The train and test focus datasets were subject to bias, given that the classes were manually defined and the foci were manually distinguished. Foci were classified one nuclear ROI at a time. Although general classes were defined, assignment of a focus’ class could deviate according to its neighboring foci in the nuclear ROI. In other words, foci would sometimes be classified with respect to its surrounding foci instead of with respect to a general definition. For this reason, the blurred class was difficult to classify by eye compared to other classes. Classification of the blurred class lowered when tested, albeit with a decent differentiation from other classes (Fig. S3, Fig. 4). Conservation of morphology was observed between datasets for the signal classes, however the distinguishing features of blurred foci were not as general. During the manual classification, we observed that blurred foci contained higher noise in the training set than the test set. Due to a one-at-a-time nuclear ROI manual focus classification approach, a discontinuity between train and test foci was observed. Future work to improve the classification of the blurred class would involve manual focus classification that classified single foci at a time, rather than a single nucleus at a time. As our method employed only two datasets, future work would incorporate 53BP1 foci from different microscopy methods in the training and testing, to increase robustness and generality of our method. Classification of a focus would then depend more on a general definition, rather than a comparison with foci in the same nucleus. Before focus classification can replace focus counting it must successfully classify cells. A future comprehensive study would include the classification of foci for the purposes of distinguishing disease as well as radiation dose, as explored by Vicar *et al* [25]. Further, data with a similar size to those obtained from an automated biodosimetry system, such as RABiT would uncover the practicality of focus classification for the purposes of distinguishing cells [52]. The detection of our foci relied on a simple pipeline (Fig. 2A-C) which could be improved upon by outsourcing the focus detection to other established software, which could automate the nuclear ROI selection and account for variation in SNR between datasets to improve focus detection without increasing the processing time.

## Conclusions

We introduce a method that classifies 53BP1 foci by quality and degree-of-overlap for 2D fluorescence microscopy images. Commonly-available machine learning methods successfully distinguished 53BP1 foci from distinct datasets using a varied parameter space. Our findings suggest the focus classes are independent of experimental conditions such as protein markers, imaging systems and protocols. Further investigation is required to determine the viability of this method for cell classification and inclusion in large-scale automated tools for radiation dose measurement. All code is publicly-available: https://github.com/mariabenjon/Fo-cusClassification.

## Abbreviations

DSB: double-stranded break
DDR: DNA Damage Response
53BP1: p-53 binding protein-1
FT: Fourier transform
KNN: k-nearest neighbors
RF: Random forest
SVM: Support vector machines

## Acknowledgements

We would like to thank Dr. Eli Rothenberg for providing the live-cell imaging setup for image acquisition of the training set and Dr. Huijun Xue for generating the EGFP-fused BP1-2 U2OS cell lines. Carsten Doil *et al* for their high content microscopy open-source data, Dr. Timothée Lionnet for key suggestions and discussions, and David Szigeti for technical support.

## Availability of data and materials

Source code and data used for the testing are included in this article. Images used for training are available from the corresponding author upon request.

## Author’s contributions

DF and MBJ conceived the project and designed experiments; DF, SJ, and MBJ defined focus classes; MBJ manually classified foci; SK and MBJ wrote the Python code; DF, SJ, and MBJ wrote the manuscript.

## Ethics approval and consent for publication

Not applicable.

## Consent for publication

Not applicable.

