## Supplementary Materials for "Machine Learning Classification of 53BP1 Foci"

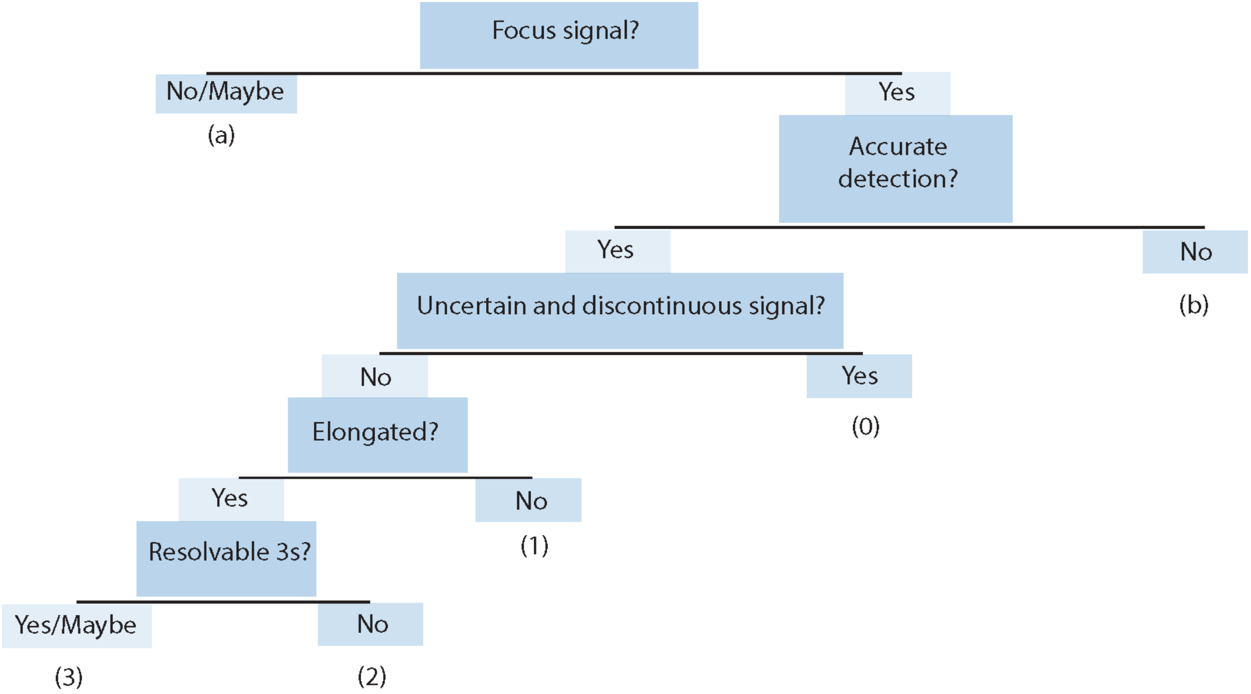


***Figure S1:*** *Manual classification decision tree. Challenging manual focus classification cases were determined with a decision tree. Classes: (a) background, (b) misdetection, (0) blurred focus, (1) focus with low overlap, (2) high overlap, and (3) moderate overlap.*

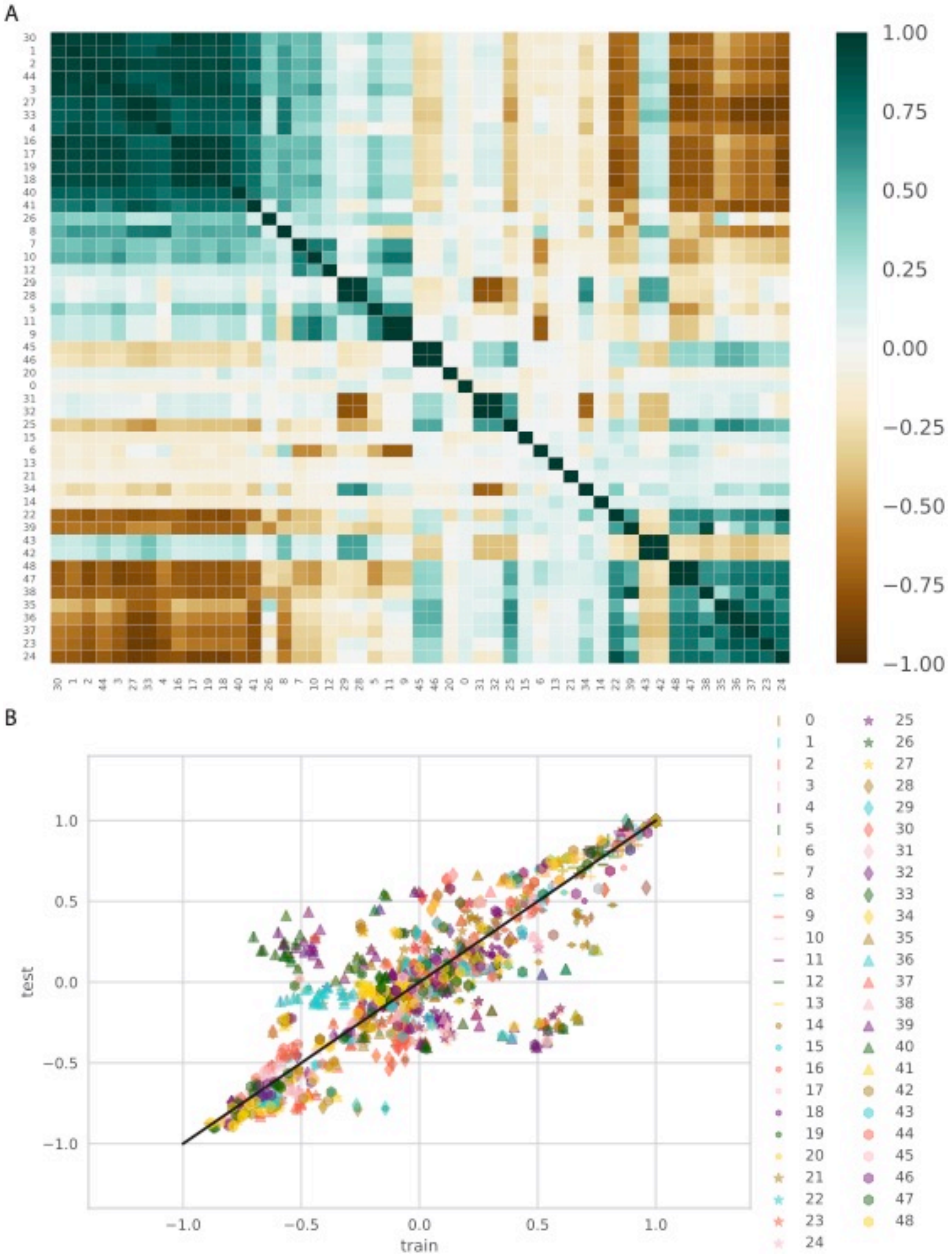
A
1.00
0.75
0.50
0.25
0.00
−0.25
−0.50
−0.75
−1.00
B
test
train
1.0
0.5
0.0
−0.5
−1.0

***Figure S2:*** *Property correlation between train and test datasets. A) Correlation heatmap of all measured properties for each detected focus and frequency spectrum-measuring variables for the test libraries. B) Scatter plot compares metric correlation between the train (Fig. 3) and test (A) sets.*

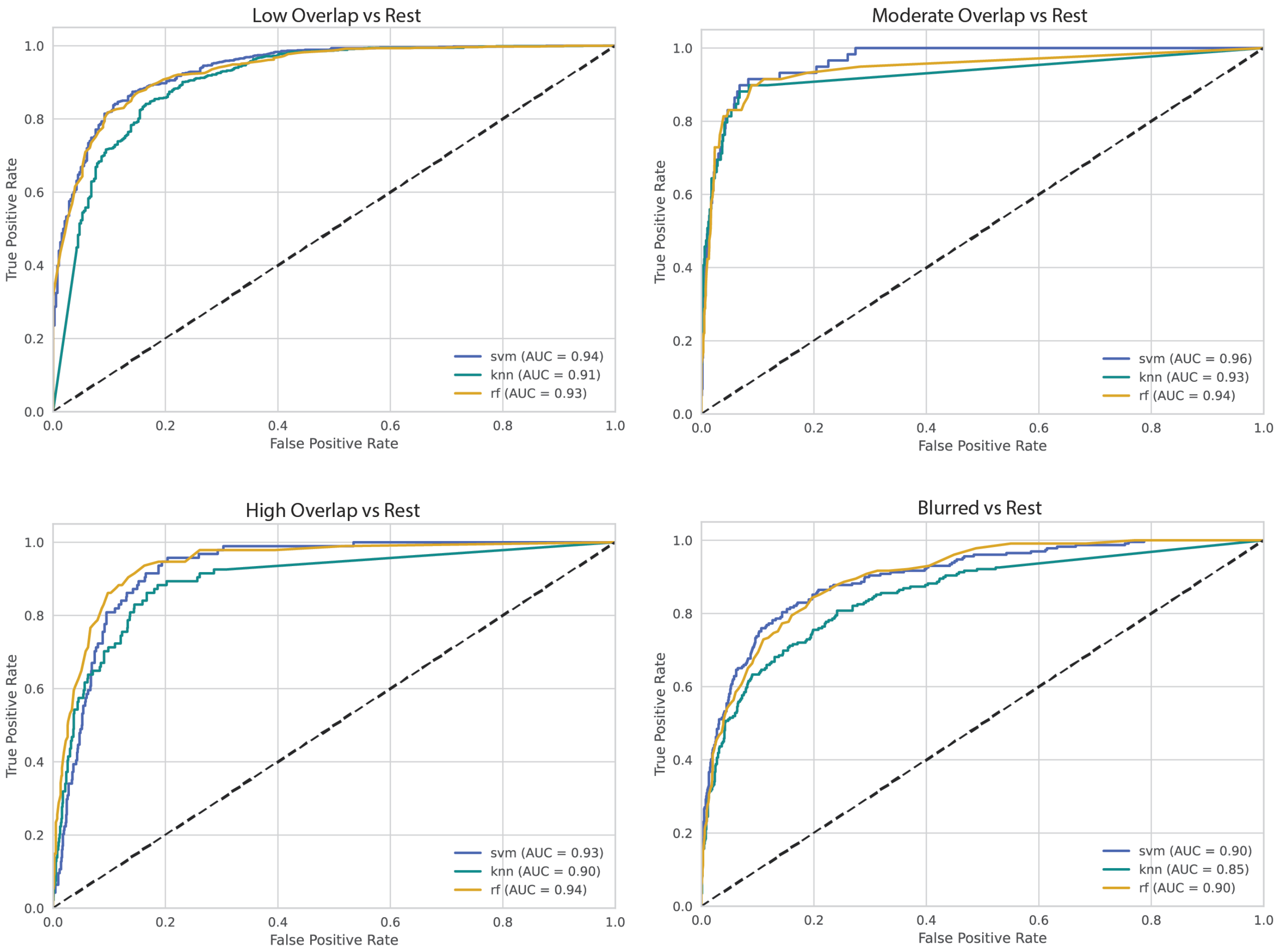


***Figure S3:*** *Classifier evaluation. Binary problems via One-vs-Rest classification measured the performance of RF, SVM, and KNN for the signal and noise classes using the train set. The ROC curves and AUC scores evaluate the classifiers' performance for each class.*

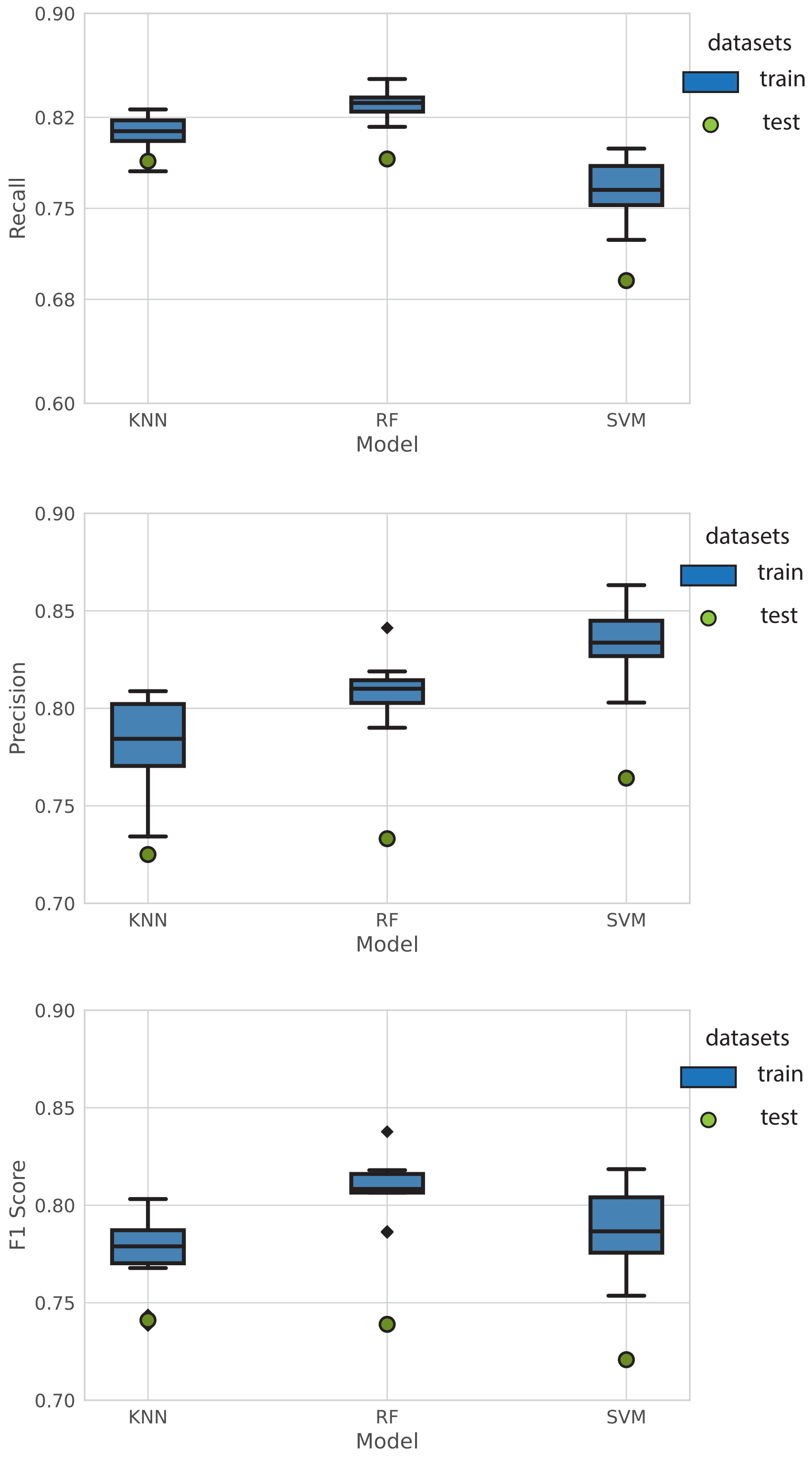

0.90
0.82
0.75
0.68
0.60
Recall
KNN
RF
SVM
Model
datasets
train
test
0.90
0.85
0.80
0.75
0.70
Precision
KNN
RF
SVM
Model
datasets
train
test
0.90
0.85
0.80
0.75
0.70
F1 Score
KNN
RF
SVM
Model
datasets
train
test

***Figure S4:*** *Multiclass problems. Ten-fold cross-validated evaluation metrics (recall, precision and F1 scores) measured the multiclass classification performance for KNN, RF, and SVM with respect to the train class (blue boxplots) and were plotted along the same classifier evaluation metrics for the test set. All metrics were weighted.*

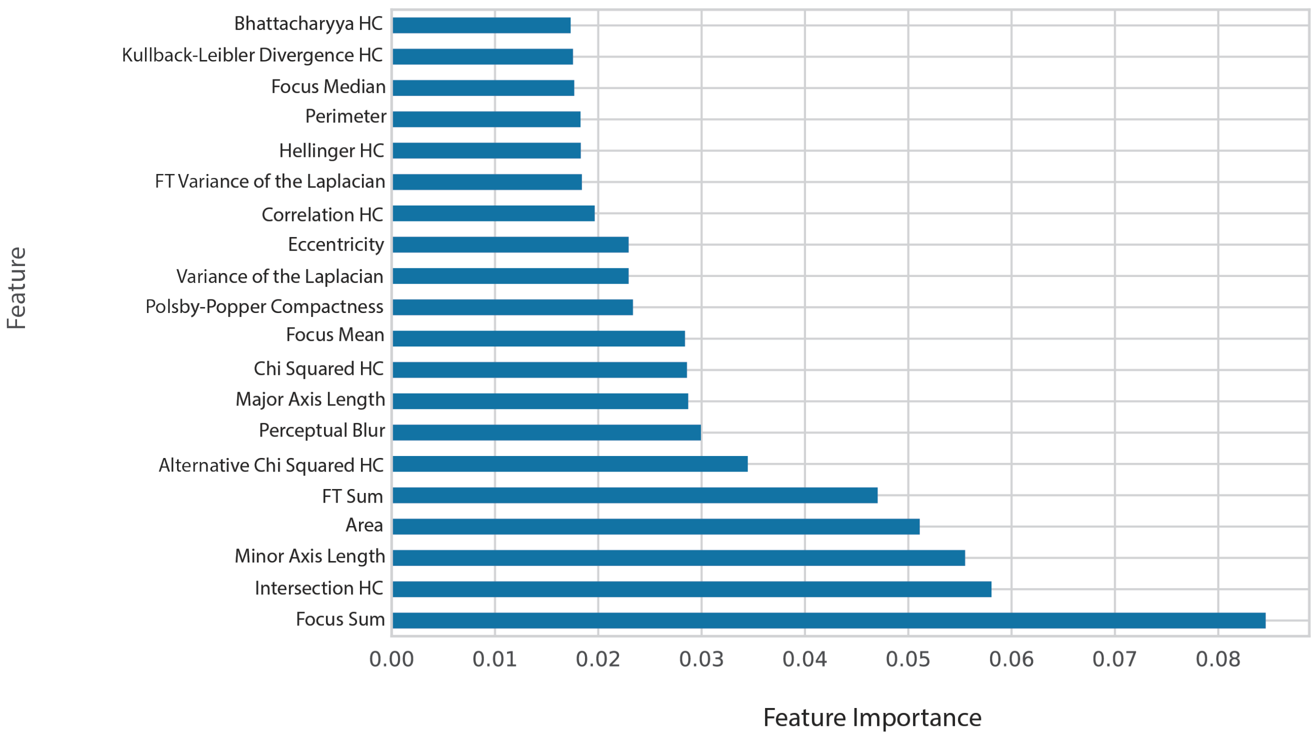


***Figure S5:*** *Feature Importance. Bar plot of top twenty most important features in Random Forest using Scikit-Learn's feature_importances_.*

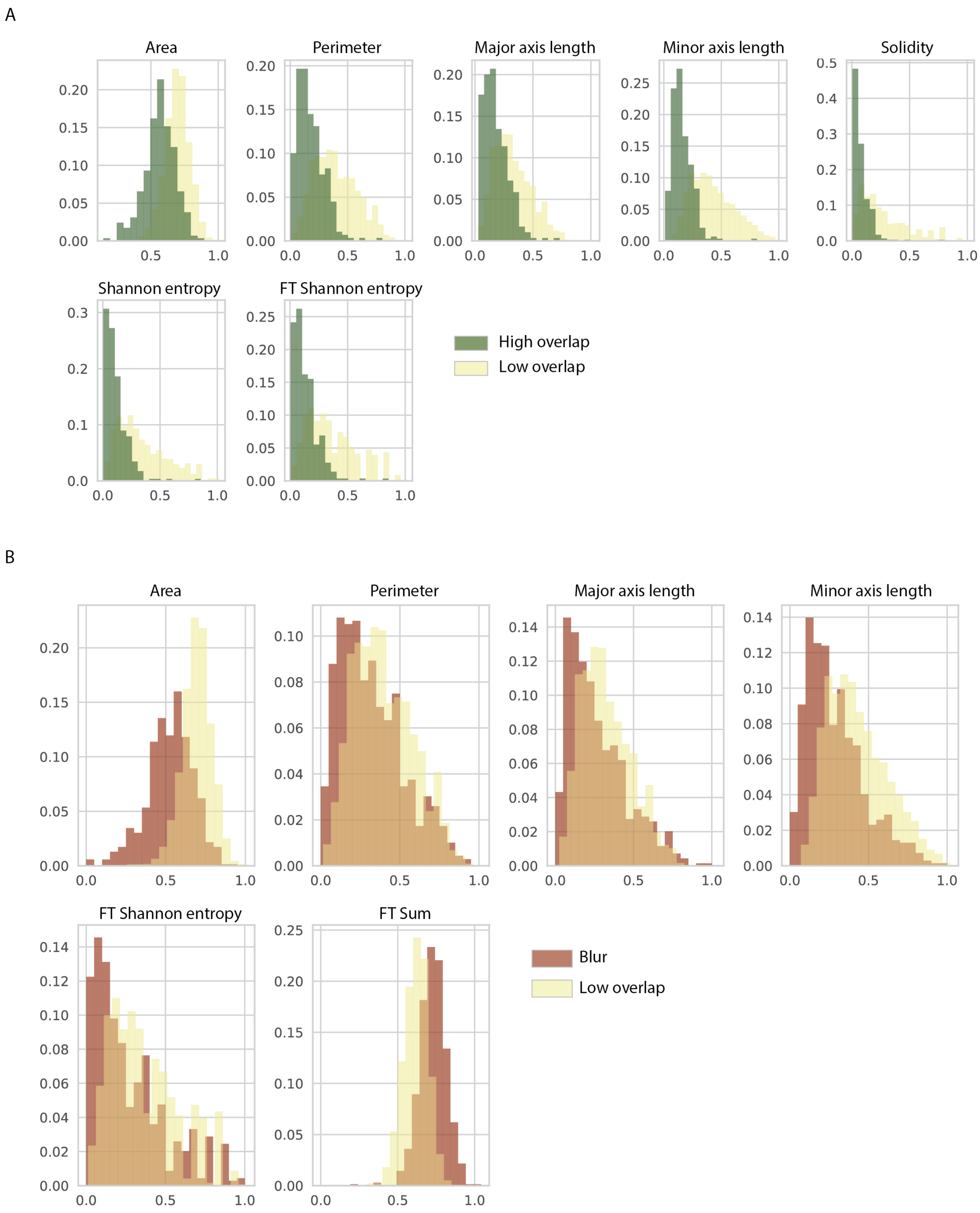


***Figure S6:*** *Focus class separation. Histograms of key properties that resolve specific focus classes. A) Symmetry metrics that separate the low overlap class (yellow) from the*

*high overlap class (green). B) Blur metrics that separate the blur class (red) from the low overlap (yellow) class.*

A

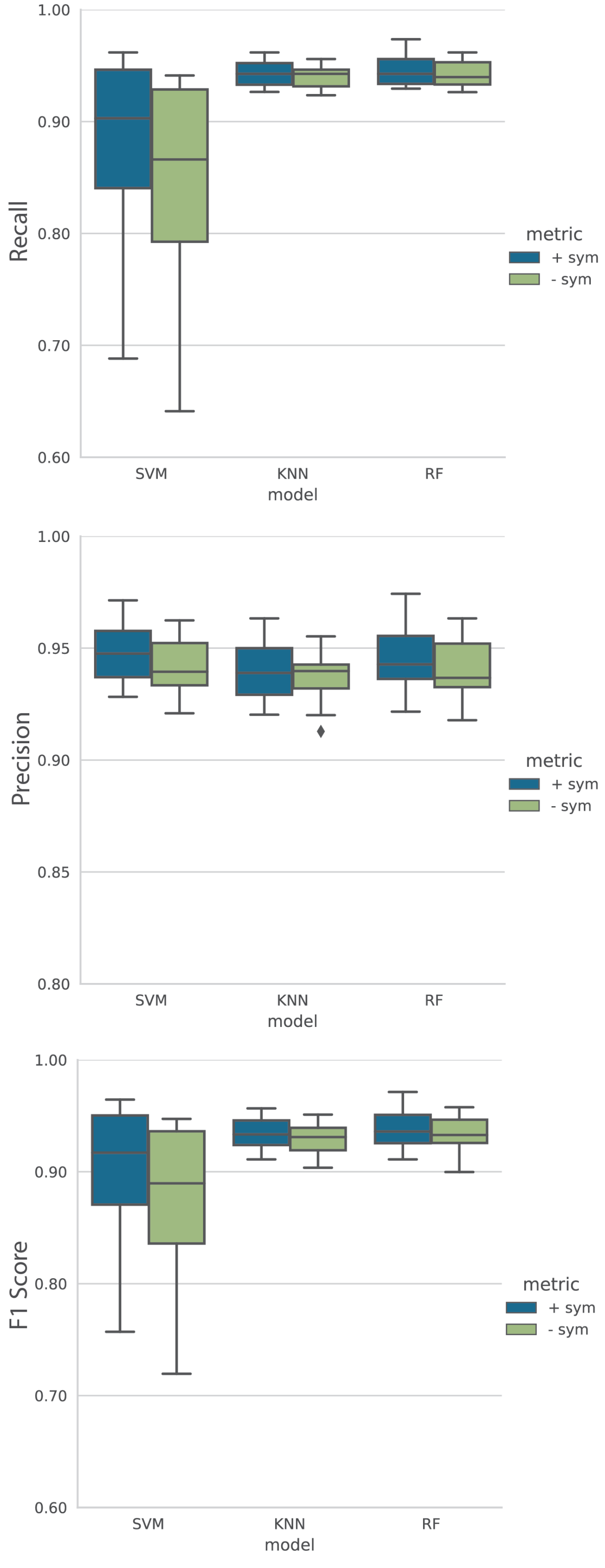
1.00
0.90
0.80
0.70
0.60
Recall
SVM
KNN
RF
model
metric
+ sym
- sym
1.00
0.95
0.90
0.85
0.80
Precision
SVM
KNN
RF
model
metric
+ sym
- sym
1.00
0.90
0.80
0.70
0.60
F1 Score
SVM
KNN
RF
model
metric
+ sym
- sym

B

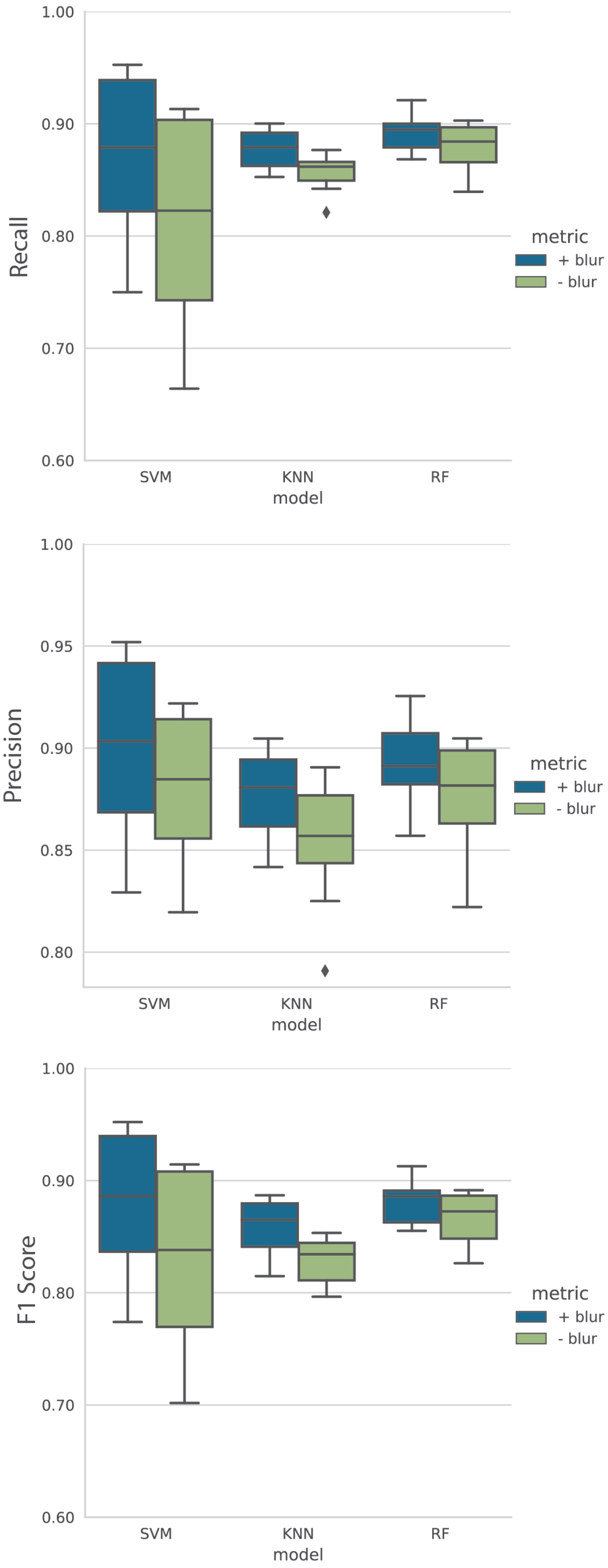
1.00
0.90
0.80
0.70
0.60
Recall
SVM
KNN
RF
model
metric
+ blur
- blur
1.00
0.95
0.90
0.85
0.80
Precision
SVM
KNN
RF
model
metric
+ blur
- blur
1.00
0.90
0.80
0.70
0.60
F1 Score
SVM
KNN
RF
model
metric
+ blur
- blur

***Figure S7:*** *Effect of metrics on classification. Boxplots of the classifiers' 10-fold cross-validated recall, precision, and F1 scores in cases where key symmetry or blur metrics were included (+, blue) and removed (-, green) to observe the effect on the differentiation between classes. A) Classification of low overlap vs high overlap class in the presence and absence of symmetry metrics for the train set. B) Distinction between blur and low overlap class in the presence and absence of blur metrics for the train sets.*
